# A gene expression correlation analysis of cell mechanosensitive systems identifies DANGER as a nuclear mechanoresponsive protein

**DOI:** 10.64898/2026.01.26.701746

**Authors:** Carmen López Morán, Lucía Garrido Amenedo, Asier Echarri

## Abstract

Cellular mechanoadaptation is a complex process involving multiple mechanotransduction pathways and mechanisms that operate in different cellular locations and organelles. Despite recent advances, the identity of all components and the molecular mechanisms of these pathways remain poorly understood. Here, we describe a strategy to identify previously unrecognized mechanotransduction components throughout the cell. Using this approach, we identify several candidate proteins involved in mechanotransduction in cellular organelles. A screen of selected candidates identified DANGER as a nuclear envelope component required for nuclear mechanoadaptation and stability. DANGER is distributed in discrete regions of the nuclear envelope. Notably, DANGER is highly enriched in bent and stretched regions of the nuclear envelope, a feature not observed for other nuclear envelope proteins associated with mechanotransduction pathways. Upon increased nuclear tension, either induced by osmotic swelling or integrin-mediated nuclear deformation, DANGER responds by forming larger clusters, suggesting that DANGER can sense changes in the nuclear envelope induced by mechanical cues. Furthermore, DANGER-depleted nuclei have a larger area, are more elongated, and are more prone to forming blebs, consistent with DANGER localizing to regions under higher tension. Together, these findings identify DANGER as a key nuclear envelope component regulating nuclear shape and nuclear envelope stability, and provide proof of concept that our gene expression correlation-based strategy can identify previously unrecognized mechanotransduction components.

## Introduction

Mechanotransduction pathways responsible for transducing mechanical signals into biochemical signals have been mostly associated with the plasma membrane ^1^, the actin cytoskeleton ^2 3^ and nucleoskeleton related components ^4,5^. In recent years, evidence has accumulated suggesting that mechanosensitive proteins, mechanotransduction pathways, and other mechanosensitive cellular components operate in many additional cellular niches ^6^. Mechanotransduction processes have been described in mitochondria ^7 8^, lysosomes ^9^, secretory pathway ^10^, endoplasmic reticulum ^11^, nuclear pore complex ^12–14^, nuclear transport ^15^, nuclear envelope ^16^ and cytoplasm related metabolic processes ^17^. This suggests that the cellular machinery involved in mechanotransduction extends well beyond canonical mechanosensory niches. Therefore, additional genes involved in sensing, transmitting and/or responding to mechanical signals are likely to exist in all these compartments, but their number and nature remain largely unknown.

Several experimental strategies have been used to identify pathways and proteins associated to mechanoadaptation, including comparative proteomic under conditions of low/high mechanical stress ^8,18^, force-dependent gene expression analysis ^19^ and biochemical approaches applied to central mechanotransduction players, such as integrins and the actin cytoskeleton^20,21^

A successful strategy to identify genes biochemically and/or functionally related to a pathway, complex, or process is based on the principle that co-expressed genes are more likely to be biochemically or functionally related than genes whose expression is weakly correlated. A seminal study provided strong evidence supporting this principle; this study showed that genes with similar expression profiles are more likely to encode interacting proteins ^22^. As expected, proteins forming permanent complexes, such as those in the ribosome, show the highest expression correlation, whereas transient complexes show less, but still significant, expression correlation ^23^. Hence, this principle has been widely used to identify and assign proteins biochemical and biological functions ^24–27^.

Cell adaptation to mechanical signals is often a broad response that includes reorganization of adhesion complexes ^28^, the actin and other cytoskeletal networks ^2,29,30^, the plasma membrane ^31^, the nucleus ^4,5^ and mitochondria ^8^, among other components ^6,32,33^. Thus, genes that encode for the main cellular components involved in sensing and transmitting mechanical cues are likely to be co-expressed with genes associated to the mechanoadaptive response in other organelles. This mechanism would facilitate a coordinated response of different cellular components to mechanical cues, consistent with the concomitant changes observed across multiple cellular components in response to the same mechanical input ^13,28,31^.

The nucleus plays a major role in adapting the cell to mechanical signals and it contains multiple components that respond to mechanical cues, including nucleoli ^34^, the chromatin ^35^, nucleoskeleton ^4^, nuclear pore complex ^13,14^ and the nuclear envelope itself ^5^. Only a handful of nuclear envelope associated proteins have been shown to respond to mechanical inputs, including the LINC complex ^36^, Emerin ^37^, cPLA2 ^16^, lamins ^4^ barrier-to-autointegration factor (BAF) ^38^ and LAP2β ^39^, representing only a small fraction of all nuclear envelope proteins. Whether there are additional nuclear envelope proteins downstream of mechanical signals remains unknown.

A proteomic study on isolated nuclear envelopes identified DANGER as a potential nuclear envelope resident ^40^. DANGER is a poorly characterized protein that was first identified as a regulator of calcium channel inositol 1,4,5-trisphosphate-receptor ^41^. In addition, several studies have linked it with death associated protein kinase (DAPK) ^42–44^, a kinase regulating apoptosis. However, the biochemical and biological functions of DANGER are still poorly understood.

Here, we describe a strategy based on gene expression correlation analysis to identify genes associated with the mechanoadaptive response of the cell. We identify all genes that positively and strongly correlate with the genes of the main mechanosensitive and mechanotransducing cellular components/pathways, i.e, focal adhesions ^45^, caveolae ^46^ and the hippo pathway ^3^. We screened some of these highly correlating genes and identified DANGER as a previously unrecognized nuclear envelope protein sensitive to mechanical cues and regulator of nuclear features impacted by mechanical stress. Thus, screening genes that strongly correlate with genes in mechanosensitive systems is useful for identifying previously unnoticed proteins downstream of mechanotransduction pathways.

## Results

### Gene expression correlation analysis of mechanotransduction systems to map organelle-associated candidate mechanotransduction components

Given that genes that are functionally related are often co-expressed ^22,23^, we hypothesized that genes co-expressing with the genes associated to the major cellular mechanotransduction pathways/structures may include genes associated with mechanotransduction pathways. As major cellular mechanotransduction pathways/structures, we selected the focal adhesion system ^45^, the Hippo pathway regulated gene set ^3^ and the major caveolae components ^46^ (Fig. 1a). These gene sets were queried against a transcriptomic data repository (see methods for details) and the top genes above the 95% percentile of positively correlating genes were selected and named the FA signature, Hippo signature and caveolar signature (Fig. 1a). All genes associated with these signatures were collectively named <u>m</u>echanotransduction-<u>c</u>orrelating genes (hereafter referred to as MC genes, Fig. 1a). In order to see how proteins encoded by these genes distributed throughout cell organelles, we crossed this list with the proteomes of cell organelles (see methods). As shown in figure 1b, the distribution of MC genes in the different organelles and cell structures was non-random; some organelles/structures contained a large fraction of MC genes, whereas others did not, suggesting that MC genes are not equally represented in cell organelles/structures. Representation of the percentage of each organelle proteome coincident with MC genes indicated that mitochondria contained very few MC genes, while the actin cytoskeleton, the plasma membrane, the nuclear envelope and the endoplasmic reticulum contained significant proportion of genes compared to other organelles (Fig. 1c). Some MC genes selected by this unbiassed strategy, and ranked in the upper part of the lists are known to be key proteins in mechanotransduction pathways. This is patent in the nuclear envelope, where lamin A/C (encoded by LMNA, ranked number 1), TMEM43 and SUN2, are all known nuclear envelope components associated with nuclear mechanotransduction ^5,47^ (Fig. 1d). In contrast, other components also showing strong correlation have not been related to mechanotransduction (for example, FNDC3B ^48^ and DANGER ^41^ in the nuclear envelope list, or MSRB3 ^49^ in the mitochondria list, Fig. 1d, e). Comparison of the three signatures indicated that there is significant overlap between them, suggesting crosstalk between the different mechanosensitive systems (Fig. 1f).

**Fig. 1.**
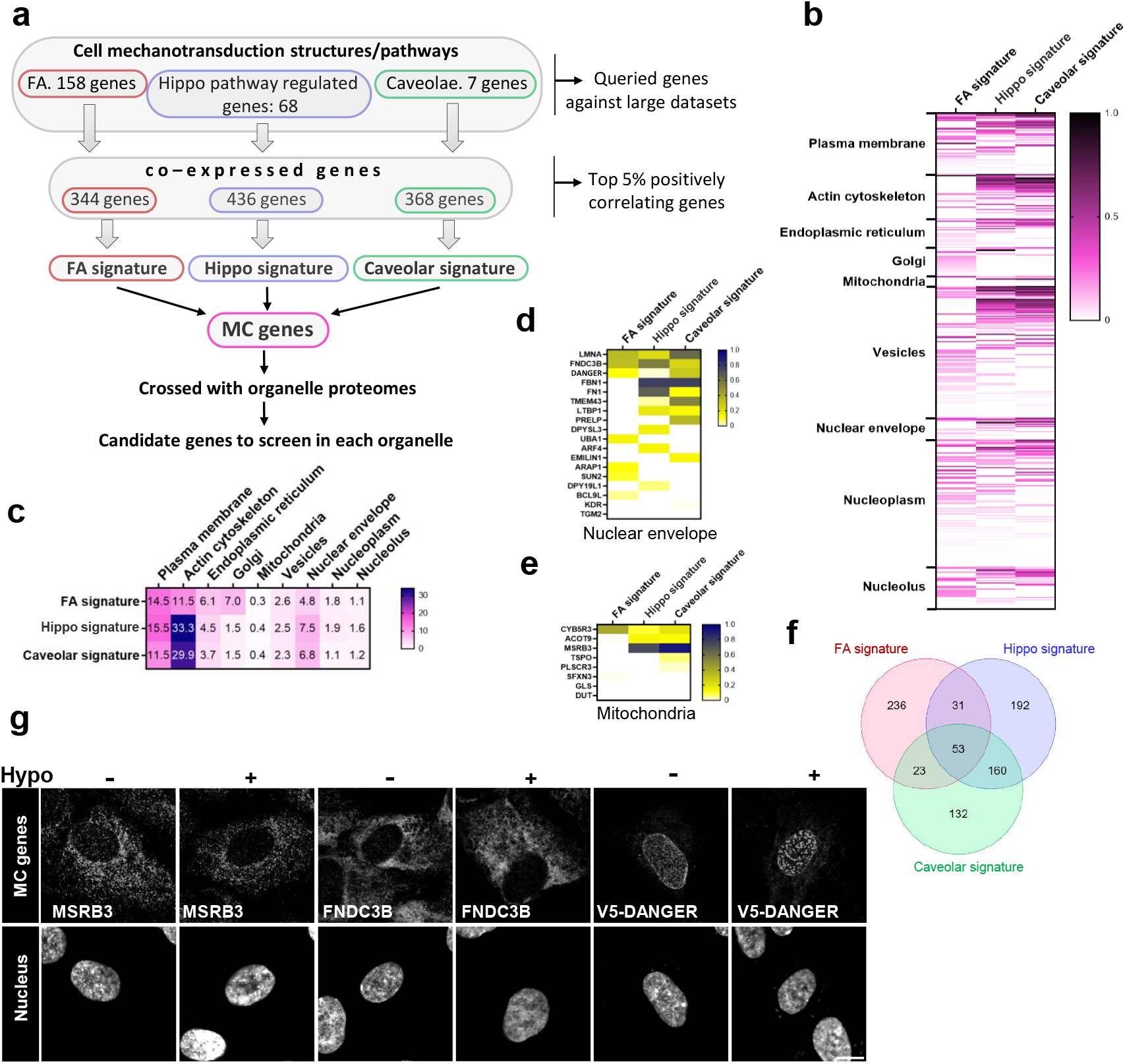
Gene expression correlation analysis identifies candidate mechanotransduction genes. a) Strategy map to identify candidate genes involved in mechanotransduction pathways. b) Heatmap of the proteins encoded by the MC genes (divided into the three signatures), ranked based on their z-score of Pearson correlation, and their presence in the different cell organelles/structures or compartments. c) Percentage of the proteins of each organelle/structure/compartment present in each of the indicated signatures. d, e) List of MC genes -separated by signatures-that are present in the nuclear envelope (d) or mitochondria (e) proteome. Proteins are ranked based on their presence in the different signatures and based on their z-score of Pearson correlation. f) Shared genes in the three signatures. g) Immunofluorescence of RPE1 cells stained for the indicated proteins and the nucleus. Cells were treated with hypo-osmotic shock and control media for 10 minutes before fixation. Scale bar 10 μm.

Thus, to test whether our strategy was useful to identify proteins associated to mechanotransduction pathways, we selected three MC genes strongly correlating with the signatures in different compartments. From the nuclear envelope we selected FNDC3B, a protein that is poorly characterized in terms of molecular function, and that has been associated with tumors ^50^; and DANGER, a protein originally isolated as a modulator of calcium signaling ^41^ (Fig. 1d). Additionally, from the mitochondria set we selected MSRB3, a member of the well-characterized methionine-R-sulfoxide reductase family, involved in regulating aging ^51,52^ (Fig. 1e).

To test empirically whether any of these proteins were associated with mechanotransduction pathways, we used a hypo-osmotic shock based assay as a means to identify proteins linked to mechanoadaptation. Hypo-osmotic conditions induce cell swelling and thereby increase tension in cell membranes, leading to the activation and/or regulation of mechanosensitive molecules resulting in a change in their localization or organization ^16,31,53^. We treated cells with iso-osmotic or hypo-osmotic medium and stained for endogenous MSRB3, FNDC3B and exogenously expressed V5-DANGER. As shown in figure 1g, we did not observe any change in MSRB3 and FNDC3B pattern, in contrast DANGER reorganized into larger clusters. Therefore, we decided to further study DANGER in the context of mechanotransduction.

### DANGER is a nuclear envelope protein enriched in stretched and highly curved regions of the nuclear envelope

DANGER is a poorly characterized protein, and its biochemical and biological functions remain largely undefined. A previous study has shown that overexpressed and tagged DANGER localizes to the nuclear envelope ^40^, confirmed in Fig. 1g. To study the function of this protein, we first validated an antibody to precisely determine the localization of its endogenous form, and to study its function in the context of mechanical signals. The signal obtained with this antibody in human cells showed a nuclear distribution (Fig. 2a). This signal was specific for DANGER, as an efficient DANGER knockdown reduced the signal in the nucleus by ≈95% (Fig. 2c; see Fig. 2b for western blot of the protein before and after silencing). This nuclear pattern was observed in different non-cancerous and cancerous cell lines (Fig. 2d), suggesting that DANGER is mostly a nuclear protein in many cell types. Overexpressed and tagged DANGER has been shown to localize in the outer nuclear envelope ^40^, to determine whether endogenous DANGER also localized to the nuclear envelope, we performed a xzy confocal acquisition and projected this stack. This analysis showed that DANGER was mostly in the periphery of the nucleus, just above DNA in the upper part of the cell (lower part in the image) and below the DNA signal in the lower part of the cell (Fig. 2e). These results are consistent with a nuclear envelope localization, both to the inner and outer nuclear membrane, although we cannot rule out that some protein may reside in the nucleoplasm.

**Fig. 2.**
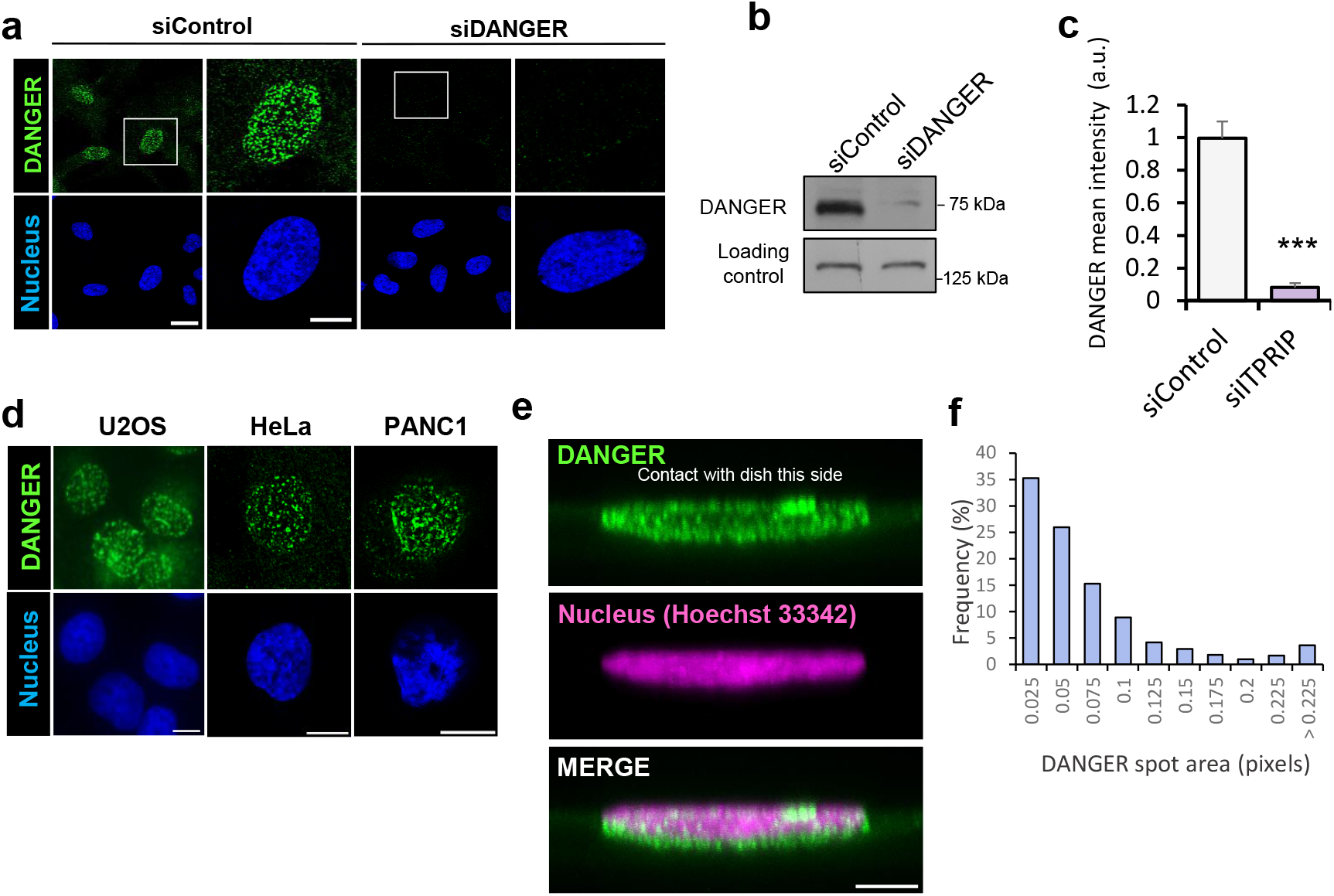
DANGER is a nuclear envelope protein forming patchy, heterogeneous populations. a) Immunofluorescence of RPE1 cells stained for DANGER and the nucleus. Cells were treated with the indicated siRNAs, fixed and stained. Scale bar 20 and 10 (inset) μm. b) Western blot of the indicated antibodies on RPE1 cell lysates transfected with the indicated siRNAs. c) Quantification of the DANGER signal in the nucleus in control and DANGER silenced RPE1 cells. Data represent mean ± s.e.m. N = 58 cells for each condition, from 3 independent experiments. d) Immunofluorescence of the indicated cells stained for the DANGER and the nucleus. Scale bar 10 μm. e) Immunofluorescence of RPE1 cells stained for the indicated proteins and the nucleus. Confocal sections were taken in the Y plane from the whole nuclear section and the maximum projection is shown. Scale bar 5 μm. f) DANGER spots are organized into heterogenous populations. Histogram of DANGER spots segmented and quantified. The frequency of each spot area range is represented. N = 1021 spots representative of 4 independent experiments.

The DANGER signal showed a patchy pattern and a non-normal size distribution skewed toward larger particles in retinal pigment epithelial cells (Fig. 2f), suggesting that DANGER organizes into heterogeneous populations. Interestingly, DANGER was markedly enriched in curved regions of the nuclear envelope, such as nuclear envelope blebs (Fig. 3a). In these blebs, DANGER was enriched approximately 5-fold compared to other nuclear envelope areas (measured in two distinct regions, Fig. 3b, c). Although these blebs were rare, about ≈70 % of all blebs contained highly enriched DANGER, suggesting that DANGER is enriched in certain types of blebs or only is recruited for a limited period. As controls of enrichment specificity, we also stained and quantified the signal for the nuclear DNA and Emerin, another nuclear envelope protein ^54^. Emerin showed a moderate enrichment in these blebs, consistent with its role in nuclear envelope stability ^54^, but much less than DANGER (Fig. 3c). In contrast, the DNA was slightly reduced (Fig. 3c). We also noticed that in multilobed nuclei, which were very rare in retinal pigment epithelial cells, DANGER was enriched in the neck connecting both lobules (not shown). To determine whether this observation was representative of DANGER or was an isolated event, we used pancreatic adenocarcinoma cell line PANC1, which has a high proportion of multilobed nuclei. In this cell line, endogenous DANGER was highly enriched in highly curved regions connecting different lobules and in stretched regions forming a bridge or neck between lobules (Fig. 3d). The enrichment in these regions was approximately 7-fold compared to other nuclear regions (Fig. 3e). To determine how specific this pattern was for a nuclear envelope protein, we stained for Emerin. Interestingly, Emerin was not enriched in these regions (Fig. 3e). Taken together, these results suggest that DANGER is a nuclear envelope protein that is highly enriched in highly curved and stretched regions of the nuclear envelope.

**Fig. 3.**
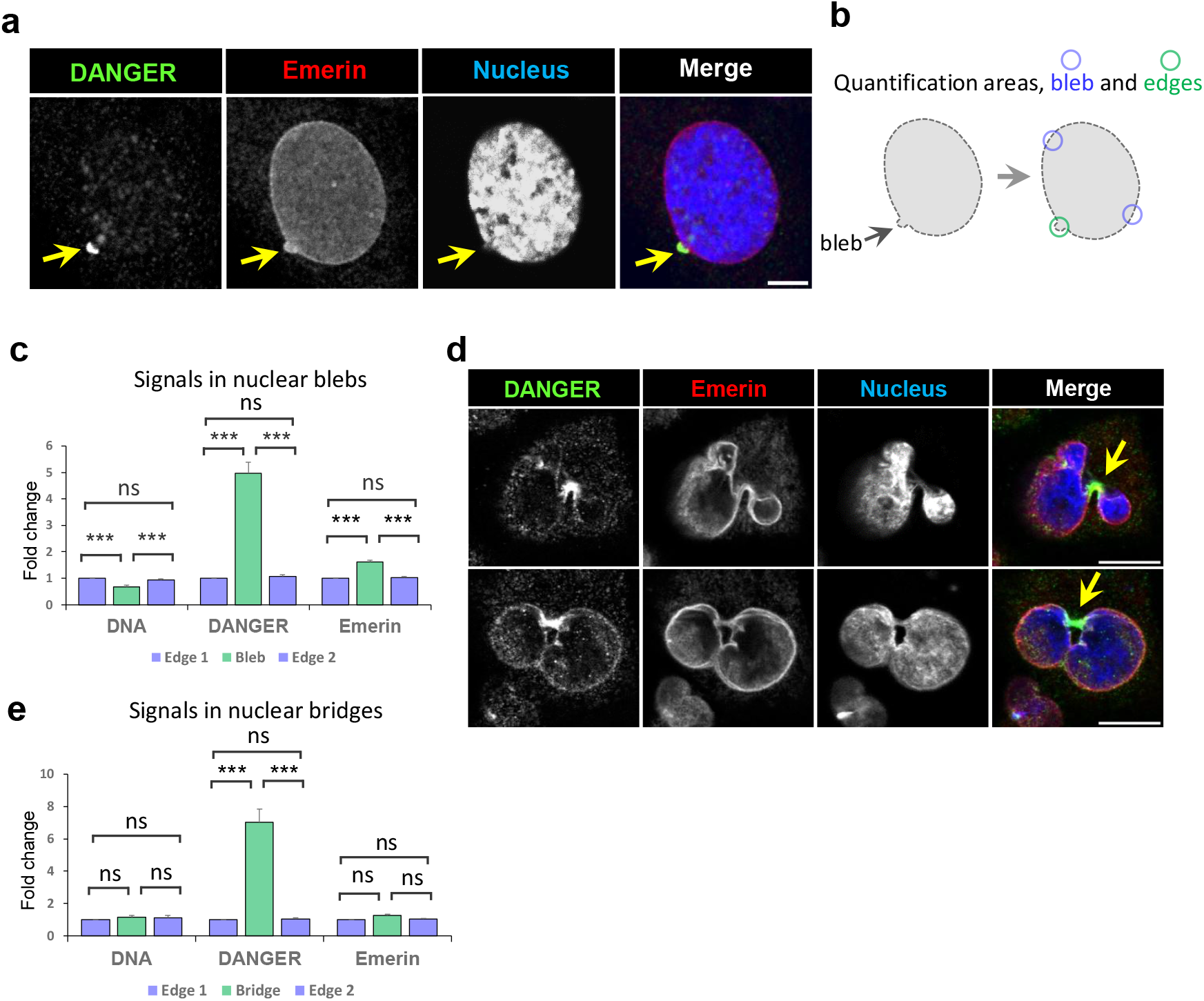
DANGER accumulates at high levels in stretched and highly curved regions of the nuclear envelope. a) Immunofluorescence of RPE1 cells stained for the indicated proteins and the nucleus. Confocal sections are shown on nuclei with blebs. The bleb is marked with an arrow. Scale bar 5 μm. b) Diagram of the three regions quantified in each nuclei containing blebs. c) Quantification of DANGER enrichment in nuclear blebs form panel a. Emerin and DNA signal was also quantified in the same regions. d) Immunofluorescence of PANC1 cells stained for the indicated proteins and the nucleus. Confocal sections are shown on multilobed nuclei with bents and bridges (marked with an arrow). Scale bar 10 μm. e) Quantification of DANGER enrichment in nuclear bridges from panel d. Emerin and DNA signal was also quantified in the same regions.

### DANGER is sensitive to increased tension in the nuclear envelope

These observations led us to hypothesize that DANGER could be sensitive to nuclear tension changes, as blebs and bridges represent a deformation of the nuclear envelope that likely increases tension locally. To test this hypothesis, we used two different approaches to increase nuclear envelope tension and subsequently analyzed the behavior of endogenous DANGER. First, we applied osmotic stress to induce nuclear swelling. As shown in figure 4a, hypo-osmotic treatment induced clustering of endogenous DANGER, similar to overexpressed and tagged DANGER (Fig. 1g). To quantify this phenotype, we segmented all DANGER spots and determined how the different DANGER spot populations changed upon nuclear envelope swelling. The median intensity of DANGER spots was significantly increased (Fig. 4b), consistent with the visual analysis (Fig. 4a). To determine what drove this increase, we next examined the full range of DANGER spots intensities and compared both treatments. We classified all detected DANGER spots in basal conditions by fluorescence intensity and divided them into four quartiles. Each quartile contained ≈25% of the spots and were labelled as: q1 (lowest intensity), q2 (low-medium), q3 (medium-high), and q4 (highest intensity) (Fig. 4c), as done before to monitor tension dependent protein reorganization ^55^. Upon osmotic swelling, DANGER spots with high intensity were significantly increased, while low intensity spots were reduced, consistent with the clustering observed in the images (Fig. 4c, a). The increase in intensity upon osmotic stress was not due to a general elevation of DANGER protein levels, as the total signal associated with the nucleus remained constant (Iso 31.6 ± 2.3, Hypo 29.0 ± 1.9, t-student test p value = 0.4). The effect of osmotic swelling was specific, as Emerin, another nuclear envelope protein, was not visibly affected (Fig. 4a).

**Fig. 4.**
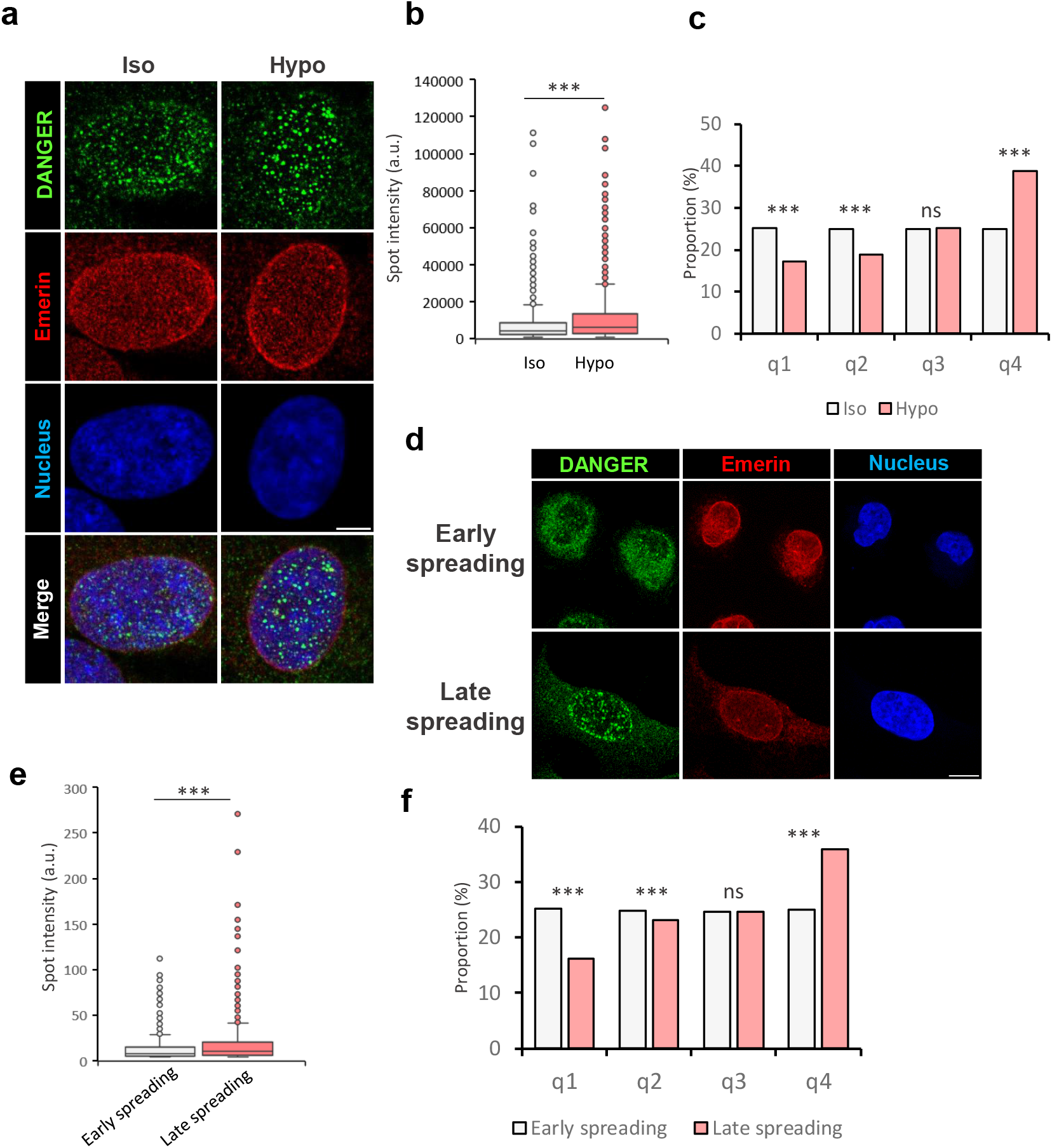
DANGER is responsive to mechanical signals. a) Immunofluorescence of RPE1 cells stained for the indicated proteins and the nucleus after treatment with iso-osmotic or hypo-osmotic medium for 10 minutes. Scale bar 20 μm. b) Quantification on DANGER spots in iso-osmotic (iso) and hypo-osmotic (Hypo) conditions. Box-and-whisker plots of spot intensity for both conditions with individual observations is shown. The box shows the IQR (25th–75th percentiles). Mann–Whitney U test p-value (two-sided) = 1.25 × 10^−17^. c) Quantification on DANGER spots intensity under iso-osmotic (iso) and hypo-osmotic (Hypo) conditions. Control spot populations (iso-osmotic) were divided into four quartiles based on spot intensity: q1 (low), q2 (low–medium), q3 (high–medium) and q4 (high). To compare proportions, a proportions test based on a χ^2^-test was used. d) Immunofluorescence of U2OS cells stained for the indicated proteins and the nucleus after spreading on a fibronectin-coated surface for 2 (early spreading) and 6 hours (late spreading). Scale bar 10 μm. e) Quantification on DANGER spots in cells spread for 2 hours (early spreading) and 6 hours (late spreading). Box-and-whisker plots of spot intensity for both conditions with individual observations is shown. The box shows the IQR (25th–75th percentiles). Mann–Whitney U test p-value (two-sided) = 1.43 × 10^−12.^. f) Quantification on DANGER spots intensity in cells spread for 2 hours (early spreading) and 6 hours (late spreading). Control spot populations (iso-osmotic) were divided into four quartiles based on spot intensity: q1 (low), q2 (low–medium), q3 (high–medium) and q4 (high). To compare proportions, a proportions test based on a χ^2^-test was used.

Osmotic swelling produces an increase in nuclear envelope tension, but also changes the ion concentration ^56^. Differentiation between these possibilities is important as DANGER has been involved in calcium signaling ^41^. In order to differentiate between ion-related effects and tension-related effects, we used a different approach to increase tension in the nucleus. When cells are platted on a FN-coated surface they spread, which induces a significant flattening and deformation of the nucleus associated with an increase in tension ^57,58^. Using this strategy, we reasoned that DANGER organization should also be sensitive to this nuclear envelope deformation. As shown in figure 4d, cell spreading induced a significant increase in nuclear area (from 171.7 ± 4.7 -2 hours-to 401.4 ± 11.2 pixels -6 hours-; t-student p value 8.9×10^-41^). Concomitantly, DANGER significantly clustered in late spreading times (6 hours) compared to early spreading (2 hours) (Fig. 4d-f). Similar to the effect of osmotic swelling, the intensity of DANGER spots increased significantly (Fig. 4e), driven by a statistically significant shift in spot populations from dim to bright spots (Fig. 4f). Taken together, these observations suggest that DANGER is sensitive to treatments that increase tension in the nuclear envelope, and that it becomes enriched in discrete regions, forming brighter clusters.

### DANGER regulates nuclear shape and nuclear envelope stability

To shed light into DANGER nuclear functions we silenced DANGER in RPE1 cells and quantified various nuclear parameters. As shown in figure 5a-c, nuclei with reduced DANGER presented a larger area and were more elongated than control siRNA silenced nuclei. To discard an off-target effect of the DANGER siRNA, we rescued the phenotype by restoring DANGER expression using an siRNA insensitive form of DANGER in DANGER silenced cells (see methods for details). This restoration, as expected, elevated the levels of DANGER (Fig. 5d), and fully rescued the nuclear area alteration to that of control cells (siControl treated cells) and the roundness decrease (Fig. 5a-c). Thus, DANGER is required to regulate the shape of the nucleus.

**Fig. 5.**
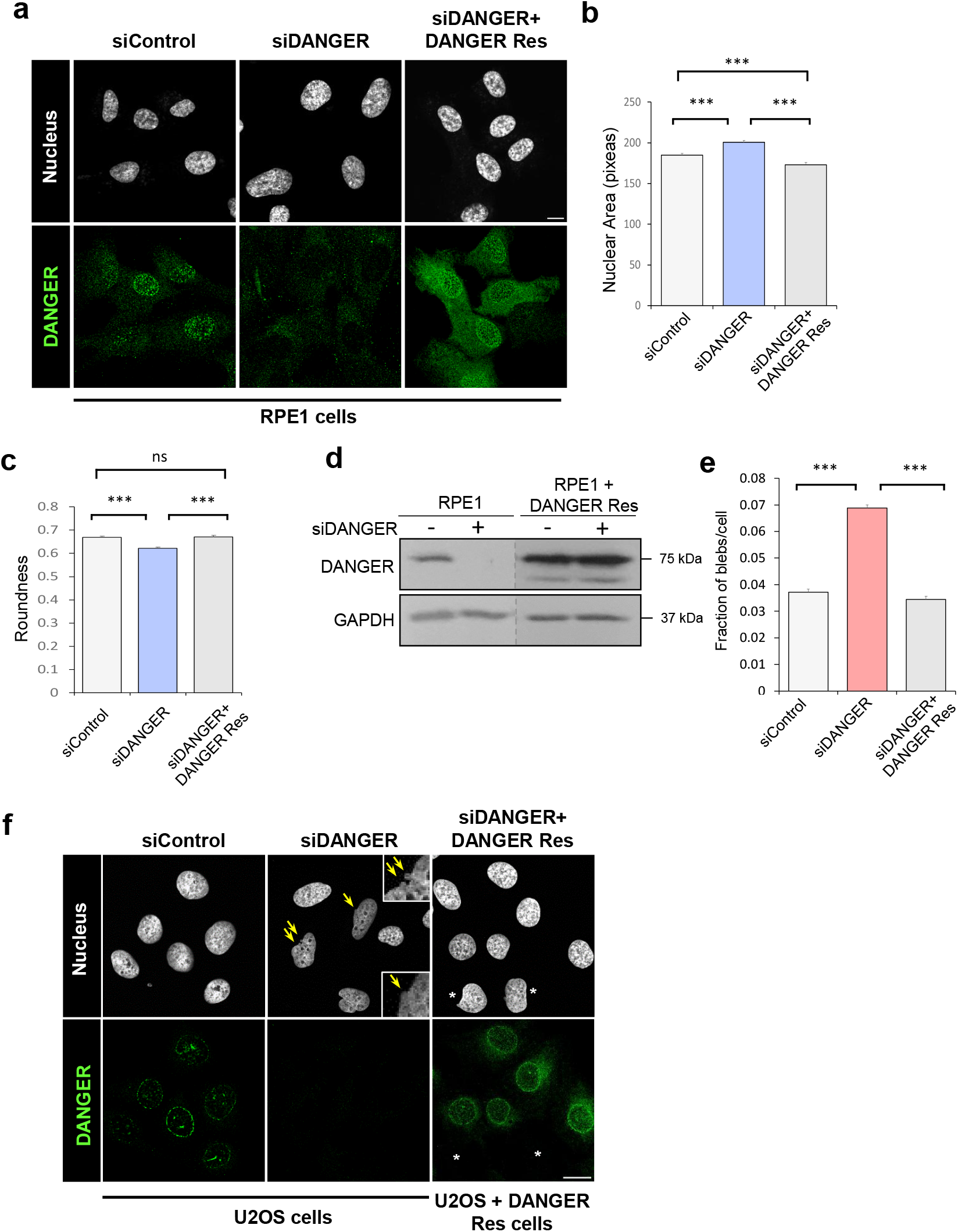
DANGER regulates nuclear shape and nuclear envelope stability. a, b, c) RPE1 cells were transfected with siRNA control (siControl), DANGER (siDANGER). siDANGER siRNA was also transfected in cells expressing siRNA insensitive DANGER cDNA (siDANGER + DANGER Res). Cells were processed for immunofluorescence, stained with Hoechst and DANGER. The indicated nuclear morphometric parameters were quantified with imageJ (b and c). The image representing overexpressed DANGER (DANGER Res) was acquired with less intensity to avoid saturation; to compare levels in this condition with respect to the endogenous see western blot below (panel d). Data represent mean ± s.e.m. Data representative of 3 independent experiments. d) Western blot showing DANGER protein levels in RPE1 cells transfected with siControl or siDANGER and in RPE1 cells expressing DANGER siRNA insensitive DANGER. A dotted line indicates the films were cut and fused. e, f) U2OS cells were transfected with siRNA control (siControl), DANGER (siDANGER). siDANGER siRNA was also transfected in U2OS cells expressing siRNA insensitive DANGER cDNA (siDANGER + DANGER Res). Cells were processed for immunofluorescence, stained with Hoechst and quantified (e) for bleb frequency in nuclei. Data represent mean ± s.e.m. N = 3040 (siControl), N=2630 (siDANGER) and N=2614 (siDANGER + DANGER Res) from 3 independent experiments. Asterisks mark two cells not re-expressing DANGER, which consistently presented nuclear deformations.

The sensitivity of DANGER to mechanical cues, its enrichment in regions of the nuclear envelope experiencing large changes in curvature (blebs) and length (elongated bridges/necks connecting nuclear lobules), and its effects on nuclear shape suggested a potential role in regulating nuclear envelope stability. To test this, we determined the role of DANGER in regulating spontaneous nuclear blebs, as these reflect nuclear envelope instability in response to mechanical stress ^59^. DANGER silenced cells had a ≈2-fold increase in the number of blebs/cell (Fig. 5e, f). This nuclear envelope instability was fully rescued by the re-expression of DANGER in DANGER silenced cells (siDANGER + DANGER Res panels, Fig. 5e, f), indicating that the effect was driven by DANGER and was not an off-target effect. It is important to mention that the exogenous DANGER expression was observed in ≈90% of cells. Meaning that some cells still had silenced DANGER, as reflected in figure 5f right panels, where two cells -marked with an asterisk-did not express DANGER, which correlated with nuclear deformations, consistent with our analysis (panel e). Collectively, these results suggest that DANGER regulates the stability of the nuclear envelope and the shape of the nucleus.

## Discussion

We developed a strategy to identify genes whose expression positively correlates with the genes associated with the main mechanotransduction systems in mammalian cells. A screening selecting three of the MC genes identified DANGER as a nuclear envelope protein acting downstream of mechanical signals. Thus, our results provide proof of concept that this strategy may be useful in future studies aimed at identifying new genes associated with mechanotransduction pathways throughout cell organelles and other regions of the cell. This strategy has been successfully used in other contexts different from mechanotransduction ^24–27^.

Cell adaptation to a chemical stimulus typically involves a sensor, transducers, and an effector; consequently, biochemical pathways often appear relatively linear ^60^. In contrast, adaptation to mechanical stimuli involves a broader response in which adhesion receptors, the cytoskeleton, the plasma membrane, cytoskeleton-associated signaling cascades, and connections to other organelles collectively drive mechanoadaptation ^61^. Therefore, some protein components across these compartments likely need to be coordinately expressed, contributing to an effective and synchronized response ^22^. Indeed, our analysis implies that there is significant coordination among the different mechanosensitive pathways based on the co-expression profiles, as there is significant overlap between the three signatures. ≈12-15% of all genes are common to the three signatures (Fig. 1f), which likely reflects the strong reciprocal functional relationship between the queried components (FA, Hippo-regulated genes and caveolae) observed empirically ^46,62–64^. These observations reinforce the idea that genes positively correlating with different mechanosensitive pathways may be part of a global mechanoadaptive cellular program, not only at the level of expression regulation, but also functionally, as proven for DANGER.

MC genes are disproportionately distributed in organelles, suggesting certain specificity and lack of randomness (Fig. 1b, c). Some large organelles, such as mitochondria had a small proportion of the selected genes, while others such as the plasma membrane and the actin cytoskeleton have a large proportion (Fig. 1b, c). Our current understanding of the abundance of mechanotransduction pathways and mechanosensitive proteins in cell organelles/structures suggests that the plasma membrane and the actin cytoskeleton contain a large number of mechanosensitive molecules and mechanotransduction pathways ^1,2,61^, which could explain our observations. An unexpected observation is the low proportion of mitochondria proteins among MC genes, even though mitochondria are known to respond strongly to mechanical cues^7,8^. This apparent discrepancy may indicate that mitochondria proteins involved in mechanoadaptation are uncoupled from the mechanosensitive systems used in this study; or alternatively, mitochondria may lack an extensive mechanotransduction network and instead rely on a small set of sensors to support mechanoadaptation.

The MC genes assigned to the nuclear envelope contained several known genes involved in mechanoadaptation ^5^, and our results suggest that DANGER is also a protein involved in mechanotransduction in the nuclear envelope. It is worth noting that the assay we used to screen for mechanoresponsive proteins measures only changes in localization in response to osmotic swelling; therefore, a negative result does not exclude involvement in mechanotransduction. Thus, FNDC3B and MSRB3 may still be relevant in mechanoadaptation. Indeed, MSRB family members directly regulate actin, an indication the they may be important for mechanoadaptation ^65^.

Moreover, our results suggest that DANGER is a nuclear envelope protein that is downstream of mechanotransduction pathways. One of the main characteristics of DANGER is its localization, which shows a heterogeneous patchy pattern, with a relatively high concentration of protein in discrete regions, compared to other nuclear envelope proteins such as Emerin ^54^. This pattern suggests that DANGER preferentially localizes to specific regions of the nuclear envelope. It is tempting to speculate that these regions correspond, at least in relative terms, to higher-tension domains, as DANGER strongly accumulates in highly bent or stretched areas, which likely represent sites of elevated membrane tension. Furthermore, DANGER tends to accumulate in discrete regions in response to osmotic swelling or cell spreading, suggesting tension-driven enrichment.

Notably, our results also indicate that DANGER is important to prevent the formation or stability of nuclear blebs. These highly bent protrusions are observed upon mechanical stress and are sites on nuclear envelope rupture ^59^, suggestive of higher tension regions. Furthermore, nuclei depleted for DANGER present a larger area, again suggestive of increased tension. Collectively, the fact that (i) DANGER is enriched in stressed regions (blebs and bridges), (ii) DANGER reorganizes in response to treatments that increase nuclear envelope tension, and (iii) it regulates nuclear parameters and features that depend on tension changes, suggest that DANGER may be an important player in modulating tension in the nuclear envelope.

### Limitations of the study

We arbitrarily selected three mechanosensitive structures/pathways (focal adhesions, caveolae, and the Hippo pathway), which, based on current knowledge, represent important cellular mechanosensing hubs. However, other mechanosensitive proteins (e.g., Piezo channels) and structures (e.g., the nuclear pore complex or nucleoli) may yield entirely different sets of correlating genes. Thus, our study may be biased toward identifying genes downstream of the selected mechanosensitive structures/pathways, and may not contain all genes involved in cell mechanoadaptation.

## Methods

### Cell culture, transfections and lentivirus production

RPE1 cells (ATCC CRL-4000) were grown in DMEM/F-12 (Lonza). HEK293 (ATCC CRL-1573), PANC1 (human pancreatic adenocarcinoma), HeLa and U2OS (human osteosarcoma, ATCC HTB-96) cells were grown in DMEM. Growth media was supplemented with 10% fetal bovine serum (FBS; Thermo Fisher Scientific), 2 mM glutamine and 100 µg/ml penicillin and streptomycin (Thermo Fisher Scientific). Cell lines were maintained in a humidified atmosphere at 37°C and 5% CO_2_. Plasmids were transfected into HEK293 cells using Fugene6 (Roche), and into RPE-1 cells using Lipofectamine 3000 (Invitrogen). siRNAs were transfected at 20 nM with RNAiMAX (Invitrogen). For lentiviral infections, HEK293 cells were infected with packaging vectors and plasmids encoding the gene of interest. All cell lines tested negative for mycoplasma.

### Reagents and plasmids

siRNA oligonucleotides were purchased from Thermofisher (silencer select). As control non-targeting siRNAs, control #2 (Dharmacon) was used. siRNA DANGER resistant to the DANGER siRNA was generated by mutating 6 targeting nucleotides (6) (GenScript). V5-DANGER was purchased from addgene. Mouse antibodies: GAPDH, V5, and Emerin. Rabbit antibodies: DANGER, FNDC3B, and MSRB3. All antibodies were used at 1:100 for IF and 1:1000 for WB.

### Immunofluorescence, microscopy and image analysis

Images were acquired on a Leica SP8 confocal microscope. ImageJ/Fiji was used to adjust brightness and contrast values. For immunostaining, cells were fixed with 4% paraformaldehyde at 37°C for 10 min, PBS washed, permeabilized with 0.2% Triton X-100 for 5 min, blocked with goat FBS, incubated with primary antibodies 1 h at room temperature, and incubated with secondary antibodies (1 h, room temperature). Nuclei were counterstained with Hoechst 33342. To detect DANGER particles, we used an ImageJ macro and particles below 9 pixels area were discarded as these were indistinguishable from the background. Individual nuclei were imaged by confocal microscopy, focusing on the basal nuclear plane. DANGER puncta/accumulations were automatically segmented using an *ad hoc* macro in Fiji (ImageJ). Rare cases in which clearly separated puncta were not properly segmented (i.e., merged into a single cluster) were excluded (≈1 out of 1000 objects). When images contained two nuclei, only the nucleus that was correctly segmented was analyzed. Nuclear area was quantified from images acquired with a 20x objective and segmented in ImageJ. All quantifications were performed blinded, whenever possible.

### Hypo-osmotic treatment

Cells were treated with 165 mOsm (hypo-osmotic condition) or 330 mOsm (iso-osmotic condition) in DMEM 10% FBS and incubated for 10 minutes at 37°C. Cells were fixed immediately without PBS washing and processed as described.

### Cell spreading on fibronectin

Cells were platted on fibronectin coated coverslips (1 μg/ml). 2 hours (early spreading) or 6 hours later cells were fixed and processed for staining.

### Generation of mechanotransduction signatures

To build the signatures associated with each mechanosensitive structure/pathway we selected all genes forming the FA (153 in total, ^45^), caveolae (7; ^46^) and those regulated by the Hippo pathway effectors YAP/TAZ (68 genes, ^3^). Each of these sets was queried against >5000 transcriptomic datasets (https://seek.princeton.edu/seek/)^66^ and the correlating genes whose z-score of Pearson correlation was above the 95th percentile of all positively correlating genes were selected (344 genes for the FA genes, 436 for the Hippo pathway regulated genes and 368 for the caveolae genes, Fig. 1a). We refer to these highly correlated gene sets as the FA signature, Hippo signature and caveolar signature. The sum of genes found in these signatures was referred to as MC genes. Since the z-scores of Pearson correlation were different for each signature, the values were normalized for comparison purposes. To cross these MC genes with the proteins of cell organelles and other structures, we used the proteomes defined in The Human Protein Atlas.

### Statistics

Statistical comparisons were carried out using two-tailed unpaired Student’s t test. At least three independent experiments are analyzed for each condition. In all figures, measurements are reported as mean ± standard error of the mean (s.e.m.), unless otherwise indicated. Mann– Whitney non-parametric test was used to compare the complete distribution of particle intensities. To compare proportions, a proportions test based on a χ^2^-test was used. P-values below or equal to 0.05, 0.01 or 0.005 were considered statistically significant and were labeled with 1, 2 or 3 asterisks respectively.

## Data availability

Data supporting the findings of this manuscript are available from the corresponding author upon reasonable request.

## Acknowledgements

We thank Alicja Tadych (Princeton University) for providing FA signature genes, and MA del Pozo for sharing reagents. This study was supported by grant I+D+i PID2022-142634NB-I00, financed by MCIN/ AEI/10.13039/501100011033/ a by FEDER Una manera de hacer Europa, granted to A.E. LGA was supported by the CSIC JAE program (JAEINT_25_01988).

## Author contribution

A.E. conceived/supervised the project, performed/analyzed experiments and wrote the manuscript. CLM and LGA designed, performed and analyzed experiments, and contributed to the writing.

## References

1. Le Roux, A.-L., Quiroga, X., Walani, N., Arroyo, M. & Roca-Cusachs, P. The plasma membrane as a mechanochemical transducer. Philos Trans R Soc Lond B Biol Sci 374, 20180221 (2019).

2. Burridge, K. & Wittchen, E. S. The tension mounts: stress fibers as force-generating mechanotransducers. J Cell Biol 200, 9–19 (2013).

3. Dupont, S. et al. Role of YAP/TAZ in mechanotransduction. Nature 474, 179–183 (2011).

4. Kirby, T. J. & Lammerding, J. Emerging views of the nucleus as a cellular mechanosensor. Nat Cell Biol 20, 373–381 (2018).

5. Echarri, A. A Multisensory Network Drives Nuclear Mechanoadaptation. Biomolecules 12, 404 (2022).

6. Phuyal, S., Romani, P., Dupont, S. & Farhan, H. Mechanobiology of organelles: illuminating their roles in mechanosensing and mechanotransduction. Trends Cell Biol 33, 1049–1061 (2023).

7. Romani, P. et al. Mitochondrial fission links ECM mechanotransduction to metabolic redox homeostasis and metastatic chemotherapy resistance. Nat Cell Biol 24, 168–180 (2022).

8. Tharp, K. M. et al. Adhesion-mediated mechanosignaling forces mitohormesis. Cell Metab 33, 1322–1341.e13 (2021).

9. Li, K. et al. Drosophila TMEM63 and mouse TMEM63A are lysosomal mechanosensory ion channels. Nat Cell Biol 26, 393–403 (2024).

10. Phuyal, S. et al. Mechanical strain stimulates COPII-dependent secretory trafficking via Rac1. EMBO J 41, e110596 (2022).

11. Rawal, S. et al. Edge curvature drives endoplasmic reticulum reorganization and dictates epithelial migration mode. Nat Cell Biol 27, 1660–1675 (2025).

12. Schuller, A. P. et al. The cellular environment shapes the nuclear pore complex architecture. Nature 598, 667–671 (2021).

13. Elosegui-Artola, A. et al. Force Triggers YAP Nuclear Entry by Regulating Transport across Nuclear Pores. Cell 171, 1397–1410.e14 (2017).

14. Zimmerli, C. E. et al. Nuclear pores dilate and constrict in cellulo. Science 374, eabd9776 (2021).

15. Andreu, I. et al. Mechanical force application to the nucleus regulates nucleocytoplasmic transport. Nat Cell Biol 24, 896–905 (2022).

16. Enyedi, B., Jelcic, M. & Niethammer, P. The Cell Nucleus Serves as a Mechanotransducer of Tissue Damage-Induced Inflammation. Cell 165, 1160–1170 (2016).

17. Park, J. S. et al. Mechanical regulation of glycolysis via cytoskeleton architecture. Nature 578, 621–626 (2020).

18. Naba, A. Ten Years of Extracellular Matrix Proteomics: Accomplishments, Challenges, and Future Perspectives. Mol Cell Proteomics 22, 100528 (2023).

19. Le, H. Q. et al. Mechanical regulation of transcription controls Polycomb-mediated gene silencing during lineage commitment. Nat Cell Biol 18, 864–875 (2016).

20. Romet-Lemonne, G. & Jégou, A. Mechanotransduction down to individual actin filaments. Eur J Cell Biol 92, 333–338 (2013).

21. Thome, S., Begandt, D., Pick, R., Salvermoser, M. & Walzog, B. Intracellular β2 integrin (CD11/CD18) interacting partners in neutrophil trafficking. Eur J Clin Invest 48 Suppl 2, e12966 (2018).

22. Ge, H., Liu, Z., Church, G. M. & Vidal, M. Correlation between transcriptome and interactome mapping data from Saccharomyces cerevisiae. Nat Genet 29, 482–486 (2001).

23. Jansen, R., Greenbaum, D. & Gerstein, M. Relating whole-genome expression data with protein-protein interactions. Genome Res 12, 37–46 (2002).

24. van Dam, S., Võsa, U., van der Graaf, A., Franke, L. & de Magalhães, J. P. Gene co-expression analysis for functional classification and gene-disease predictions. Brief Bioinform 19, 575–592 (2018).

25. Marcotte, E. M., Pellegrini, M., Thompson, M. J., Yeates, T. O. & Eisenberg, D. A combined algorithm for genome-wide prediction of protein function. Nature 402, 83–86 (1999).

26. Pujana, M. A. et al. Network modeling links breast cancer susceptibility and centrosome dysfunction. Nat Genet 39, 1338–1349 (2007).

27. Kolberg, L., Kerimov, N., Peterson, H. & Alasoo, K. Co-expression analysis reveals interpretable gene modules controlled by trans-acting genetic variants. Elife 9, e58705 (2020).

28. Kechagia, J. Z., Ivaska, J. & Roca-Cusachs, P. Integrins as biomechanical sensors of the microenvironment. Nat Rev Mol Cell Biol 20, 457–473 (2019).

29. Alisafaei, F. et al. Vimentin is a key regulator of cell mechanosensing through opposite actions on actomyosin and microtubule networks. Commun Biol 7, 658 (2024).

30. Li, Y. et al. Compressive forces stabilize microtubules in living cells. Nat Mater 22, 913–924 (2023).

31. Echarri, A. et al. An Abl-FBP17 mechanosensing system couples local plasma membrane curvature and stress fiber remodeling during mechanoadaptation. Nat Commun 10, 5828 (2019).

32. Kai, F., Leidal, A. M. & Weaver, V. M. Tension-induced organelle stress: an emerging target in fibrosis. Trends Pharmacol Sci 46, 117–131 (2025).

33. Goodman, M. B., Haswell, E. S. & Vásquez, V. Mechanosensitive membrane proteins: Usual and unusual suspects in mediating mechanotransduction. J Gen Physiol 155, e202213248 (2023).

34. Pundel, O. J., Blowes, L. M. & Connelly, J. T. Extracellular Adhesive Cues Physically Define Nucleolar Structure and Function. Adv Sci (Weinh) 9, e2105545 (2022).

35. Tajik, A. et al. Transcription upregulation via force-induced direct stretching of chromatin. Nat Mater 15, 1287–1296 (2016).

36. Arsenovic, P. T. et al. Nesprin-2G, a Component of the Nuclear LINC Complex, Is Subject to Myosin-Dependent Tension. Biophys J 110, 34–43 (2016).

37. Henretta, S. & Lammerding, J. Nuclear envelope proteins, mechanotransduction, and their contribution to breast cancer progression. NPJ Biol Phys Mech 2, 14 (2025).

38. Halfmann, C. T. et al. Repair of nuclear ruptures requires barrier-to-autointegration factor. J Cell Biol 218, 2136–2149 (2019).

39. Sun, J. et al. LAP2β transmits force to upregulate genes via chromatin domain stretching but not compression. Acta Biomater 163, 326–338 (2023).

40. Cheng, Q. et al. Quantitative Proteomics Reveals Significant Downregulation of Glutathione Metabolism in Sepsis-Induced Liver Injury. J Proteome Res https://doi.org/10.1021/acs.jproteome.5c00912 (2026) xdoi:10.1021/acs.jproteome.5c00912.

41. van Rossum, D. B. et al. DANGER, a novel regulatory protein of inositol 1,4,5-trisphosphate-receptor activity. J Biol Chem 281, 37111–37116 (2006).

42. Cao, C. et al. ITPRIP promotes glioma progression by linking MYL9 to DAPK1 inhibition. Cell Signal 85, 110062 (2021).

43. Kwon, T. et al. DANGER is involved in high glucose-induced radioresistance through inhibiting DAPK-mediated anoikis in non-small cell lung cancer. Oncotarget 7, 7193–7206 (2016).

44. Kang, B. N. et al. Death-associated protein kinase-mediated cell death modulated by interaction with DANGER. J Neurosci 30, 93–98 (2010).

45. Zaidel-Bar, R., Itzkovitz, S., Ma’ayan, A., Iyengar, R. & Geiger, B. Functional atlas of the integrin adhesome. Nat Cell Biol 9, 858–867 (2007).

46. Echarri, A. & Del Pozo, M. A. Caveolae - mechanosensitive membrane invaginations linked to actin filaments. J Cell Sci 128, 2747–2758 (2015).

47. Liang, W.-C. et al. TMEM43 mutations in Emery-Dreifuss muscular dystrophy-related myopathy. Ann Neurol 69, 1005–1013 (2011).

48. Jiang, H., Chu, B. L., He, J., Liu, Z. & Yang, L. Expression and prognosis analyses of the fibronectin type-III domain-containing (FNDC) protein family in human cancers: A Review. Medicine (Baltimore) 101, e31854 (2022).

49. Kim, H.-Y. & Gladyshev, V. N. Methionine sulfoxide reduction in mammals: characterization of methionine-R-sulfoxide reductases. Mol Biol Cell 15, 1055–1064 (2004).

50. Jiang, H., Chu, B. L., He, J., Liu, Z. & Yang, L. Expression and prognosis analyses of the fibronectin type-III domain-containing (FNDC) protein family in human cancers: A Review. Medicine (Baltimore) 101, e31854 (2022).

51. Kim, H.-Y. & Gladyshev, V. N. Methionine sulfoxide reduction in mammals: characterization of methionine-R-sulfoxide reductases. Mol Biol Cell 15, 1055–1064 (2004).

52. Koc, A. & Gladyshev, V. N. Methionine sulfoxide reduction and the aging process. Ann N Y Acad Sci 1100, 383–386 (2007).

53. Sinha, B. et al. Cells respond to mechanical stress by rapid disassembly of caveolae. Cell 144, 402–413 (2011).

54. Lammerding, J. et al. Abnormal nuclear shape and impaired mechanotransduction in emerin-deficient cells. J Cell Biol 170, 781–791 (2005).

55. Echarri, A. et al. Caveolar domain organization and trafficking is regulated by Abl kinases and mDia1. J Cell Sci 125, 3097–3113 (2012).

56. Finan, J. D. & Guilak, F. The effects of osmotic stress on the structure and function of the cell nucleus. J Cell Biochem 109, 460–467 (2010).

57. Colbert, M.-J., Raegen, A. N., Fradin, C. & Dalnoki-Veress, K. Adhesion and membrane tension of single vesicles and living cells using a micropipette-based technique. Eur Phys J E Soft Matter 30, 117–121 (2009).

58. Khatau, S. B. et al. A perinuclear actin cap regulates nuclear shape. Proc Natl Acad Sci U S A 106, 19017–19022 (2009).

59. Clark, W. H. Electron microscope studies of nuclear extrusions in pancreatic acinar cells of the rat. J Biophys Biochem Cytol 7, 345–352 (1960).

60. Jordan, J. D., Landau, E. M. & Iyengar, R. Signaling networks: the origins of cellular multitasking. Cell 103, 193–200 (2000).

61. Iskratsch, T., Wolfenson, H. & Sheetz, M. P. Appreciating force and shape—the rise of mechanotransduction in cell biology. Nat Rev Mol Cell Biol 15, 825–833 (2014).

62. Wang, L. et al. Integrin-YAP/TAZ-JNK cascade mediates atheroprotective effect of unidirectional shear flow. Nature 540, 579–582 (2016).

63. Rausch, V. & Hansen, C. G. The Hippo Pathway, YAP/TAZ, and the Plasma Membrane. Trends Cell Biol 30, 32–48 (2020).

64. Nardone, G. et al. YAP regulates cell mechanics by controlling focal adhesion assembly. Nat Commun 8, 15321 (2017).

65. Lee, B. C. et al. MsrB1 and MICALs regulate actin assembly and macrophage function via reversible stereoselective methionine oxidation. Mol Cell 51, 397–404 (2013).

66. Zhu, Q. et al. Targeted exploration and analysis of large cross-platform human transcriptomic compendia. Nat Methods 12, 211–214, 3 p following 214 (2015).

